# Generalized Hidden Markov Models for Phylogenetic Comparative Datasets

**DOI:** 10.1101/2020.07.18.209874

**Authors:** James D. Boyko, Jeremy M. Beaulieu

**Affiliations:** Department of Biological Sciences, University of Arkansas, Fayetteville, AR, 72701, USA

**Keywords:** ancestral state reconstruction, discrete character evolution, hidden Markov models, hidden states, stochastic map

## Abstract

1. Hidden Markov models (HMM) have emerged as an important tool for understanding the evolution of characters that take on discrete states. Their flexibility and biological sensibility make them appealing for many phylogenetic comparative applications.
2. Previously available packages placed unnecessary limits on the number of observed and hidden states that can be considered when estimating transition rates and inferring ancestral states on a phylogeny.
3. To address these issues, we expanded the capabilities of the R package `corHMM` to handle *n*-state and *n*-character problems and provide users with a streamlined set of functions to create custom HMMs for any biological question of arbitrary complexity.
4. We show that increasing the number of observed states increases the accuracy of ancestral state reconstruction. We also explore the conditions for when an HMM is most effective, finding that an HMM is an appropriate model when the degree of rate heterogeneity is moderate to high.
5. Finally, we demonstrate the importance of these generalizations by reconstructing the phyllotaxy of the ancestral angiosperm flower. Partially contradicting previous results, we find the most likely state to be a whorled perianth, whorled androecium, whorled gynoecium. The difference between our analysis and previous studies was that our modeling explicitly allowed for the correlated evolution of several flower characters.

## 1. Introduction

Hidden Markov models (HMMs) are important centerpieces in many biological applications (Eddy, 2004; Yang Lou, 2017). They provide a natural framework for comparative biologists, particularly for relaxing assumptions about homogeneous evolution through time and across taxa without vastly increasing the number of parameters (e.g., Felsenstein & Churchill, 1996; Galtier, 2001; Penny, McComish, Charleston, & Hendy, 2001; Beaulieu, O’Meara, & Donoghue, 2013; Beaulieu & O’Meara, 2016). For instance, simple models of binary character evolution make sense for small, young clades, because a single set of transition rates seems like a reasonable assumption. However, homogeneous rates are unlikely to explain the evolution of the same character across a much larger and older clade in which transition rates may differ dramatically among subclades, perhaps due to correlations with traits that were not included in the model. This observation was the motivation for the development of the hidden rate model (HRM) of Beaulieu *et al*. (2013), which uses a hidden Markov approach to objectively locate regions of a phylogeny where hidden factors have either promoted or constrained the evolutionary process for a binary character.

Within comparative biology, HMMs have been applied as both standalone models (Beaulieu et al., 2013) and in combination with other phylogenetic models (e.g., hidden state-dependent speciation and extinction models, Beaulieu & O’Meara, 2016). Hidden Markov models can be used to address many problems in comparative biology (Siepel & Haussler, 2005) and their flexibility allows biologists to create models tailored to their specific hypotheses. However, previous implementations of HMMs for comparative methods have placed limitations on the number of observed and hidden states. For instance, the implementation of the HRM model of Beaulieu et al. (2013) is restricted only to the analysis of binary characters. There is no mathematical basis for limiting the number of observed states or hidden states in an HMM, and such constraints necessitate a simplification of datasets and candidate models.

Here we describe a new version of `corHMM` that implements *n*-state HMMs. This does not require new algorithms or a different likelihood function. Instead, we optimized and generalized existing code so users can create custom HMMs for any biological question of arbitrary complexity. We have also added a number of “quality of life” improvements that make `corHMM` much easier to use and interpret, including an implementation of stochastic character mapping (simmap; Bollback, 2006). Additionally, we demonstrate the effectiveness of HMMs to identify rate heterogeneity when it is present, and we outline the informational advantages of increasing the number of observed and hidden states in discrete character data sets. Finally, to demonstrate the importance of this generalization, we apply `corHMM` to reconstruct the morphology of the ancestral angiosperm flower.

## 2. Materials and Methods

### 2.1 Generalizing HMMs

From a technical standpoint, hidden Markov models have a hierarchical structure that can be broken down into two components: a “state-dependent process” (Fig. 1a,b) and an unobserved “parameter process” (Fig. 1c)(Zucchini, MacDonald, & Langrock, 2017). In comparative biology, for characters that take on discrete states the standard “state-dependent process” is a continuous-time Markov chain with finite state-space (CTMC-FS). The benefit of a Markov model is its simplicity — to calculate the probabilities of observed discrete states at the tips of a phylogeny all that is required is a tree, a transition model describing transitions among a set of observed states, and frequencies at the root (O’Meara, 2012; Fig. 1a,b). The observed states could be any discretized trait such as presence or absence of extrafloral nectaries (Marazzi et al., 2012), woody or herbaceous growth habit (Beaulieu et al., 2013), or diet state across all animals (Román-Palacios, Scholl, & Wiens, 2019). However, a simple Markov process that assumes homogeneity through time and across taxa is often not adequate to capture the variation of real datasets (e.g. Beaulieu *et al*., 2013). Under an HMM, observations are generated by a given state-dependent process, which in turn depends on the state of the parameter process. In other words, the observed data are the product of several processes occurring in different parts of a phylogeny and the parameter process is way of linking them. It is initially unknown what the parameter process corresponds to biologically, hence the moniker “hidden” state. Nevertheless, the information for detecting hidden states comes from the differences in how the observed states change. As long as the transitions between observed states of different lineages are more adequately described by several Markov processes rather than a single process, there will be information to detect hidden states (see *3.1 Performance in Simulations*).

**Figure 1.**
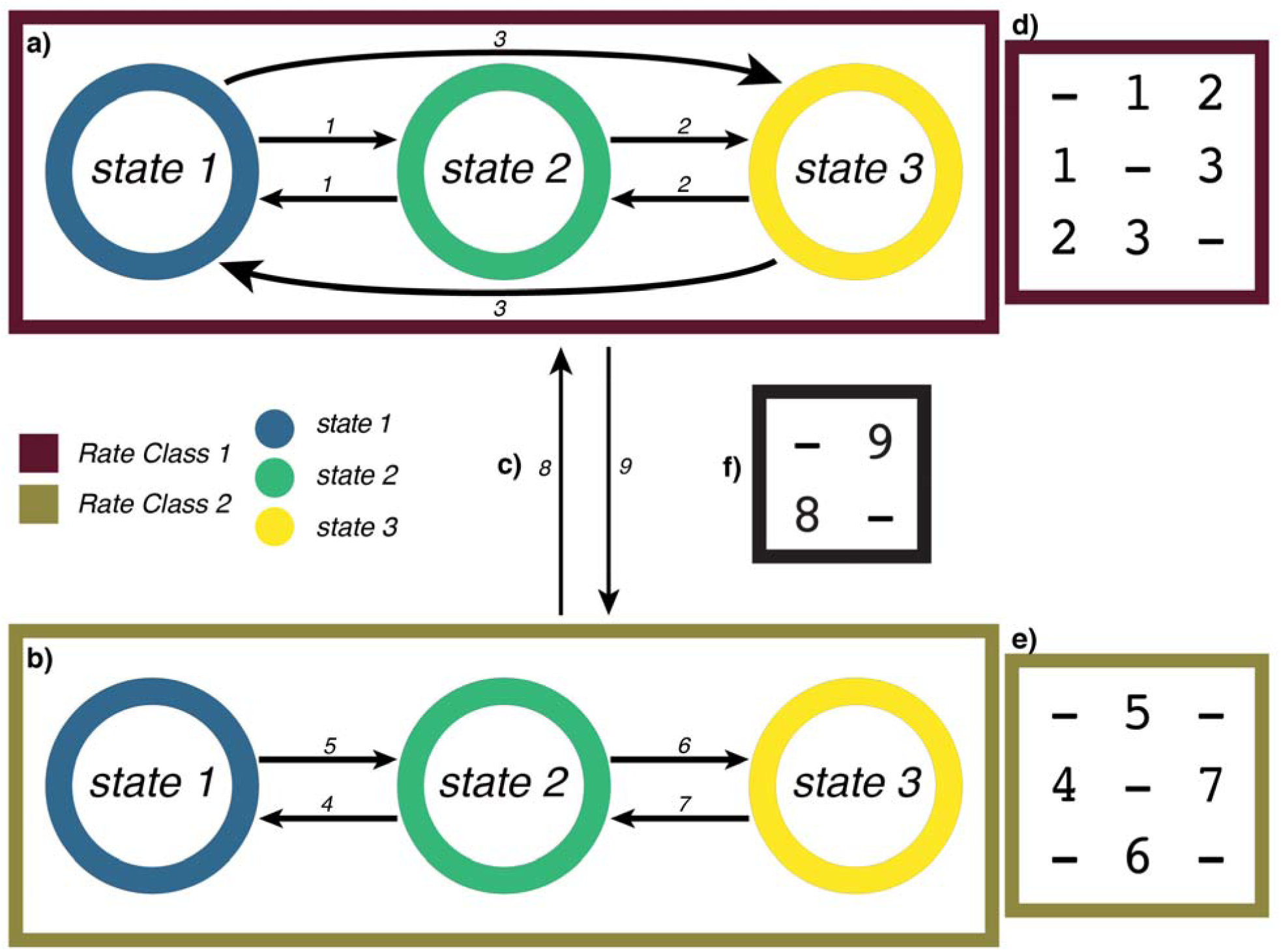
A decomposed HMM containing 3 observed states and 2 hidden rate classes. R1 is one state-dependent process that describes transitions to and from observed states as being equal (a, d), whereas R2 is a state-dependent process that describes state 2 as a necessary intermediate (b, e). The parameter process that relates R1 and R2 and describes the transitions between R1 and R2 (c, f).

The likelihood of any HMM is obtained by maximizing the standard likelihood formula, *L* = *P*(*D*|**Q**,*T*), for observing character states, *D*, across a set of extant taxa, given the continuous-time Markov model **Q**, and a fixed topology with a set of branch lengths (denoted by *T*). For a binary character, **Q** is a 2×2 transition matrix representing the transition rates, whose entries define transitions between the character states, 0 and 1. To form an HMM, we expand **Q** to accommodate both observed and hidden states. Formally, the HMM can be generalized to include any number of observed states (e.g., 0, 1, 2), and hidden states (e.g., A, B, C). Following Beaulieu and O’Meara (2016), the state space is defined as *o* being the index of the observed state, *o* ∈ 0,1,…,*α*, and *h* as the index of the hidden state, *h* ∈ *A,B*,…,*β*. Thus, a given model will have, in general, |*o*|,|*h*| states. In `corHMM`, the model **Q** is defined by amalgamating each of the state-dependent processes and the parameter process specified in the model. For example, if we have state-dependent matrices, ***R***,

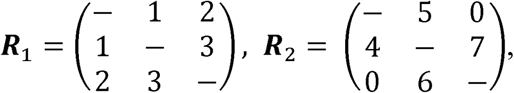

that are related by a parameter-process ***P***,

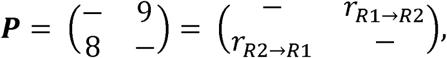

where entries *r*_R1→R2_ and *r*_R2→R1_ define transition rates between the state-dependent processes, we can extend Eq. 2 of Tarasov (2019) to amalgamate these processes,

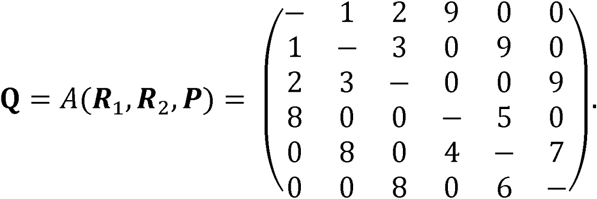

This matrix can be understood as a block matrix where the diagonal blocks are the state-dependent processes ***R***_1_ and ***R***_2_, and the off-diagonal blocks are the parameter process, ***P***, that describe transitions between ***R***_1_ and ***R***_2_. It is important to note that although we amalgamate the matrices above, it is still possible for users to modify the transitions between hidden states.

The general formulation of an HMM can easily be extended to examine the correlated evolution of multiple characters (Pagel, 1994). For example, consider a case of two binary characters where trait 1 defines the presence or absence of fleshiness of fruits, and trait 2 defines whether or not the fruits are animal-dispersed. At most there are four binary combinations of these characters (i.e., 00, 01, 10, and 11). But, it can also be coded as a single multistate character, where 1=dry fruits not dispersed by animals, 2=dry fruits dispersed by animals, 3=fleshy fruits not dispersed by animals, and 4=fleshy fruits dispersed by animals. Therefore, transforming binary combinations to multistate characters also applies for two characters with a different number of observed states. In other words, one character could be binary (e.g., dry vs. fleshy fruit) and the other could be multistate (e.g., fruits dispersed mechanically, by wind, or by animal).

### 2.2 Simulation Study

We conducted a set of simulations to address two broad goals. First we tested whether there is an informational advantage to increasing the number of observed states by comparing two-state, three-state, and four-state datasets. Our second goal was to test the ability of hidden Markov models to detect varying degrees of rate heterogeneity. We then linked these goals together by testing whether HMMs can recover some of the informational content of unobserved characters through the use of hidden states. These simulations are in no way exhaustive, but represent a set of reasonable questions that many beginning users might have about the behavior of HMMs.

#### 2.2.1 Increasing the number of observed characters or states

To test the behavior of two-state, three-state, and four-state datasets we relied on ancestral state reconstruction (ASR) at nodes. ASR is a widely-utilized feature of `corHMM`, and it is important to know the accuracy of multistate ancestral reconstructions. Additionally, using ancestral states gives us a direct means to compare models with different datasets. A 250-tip phylogeny was simulated (birth rate set to 1 event Myr^-1^, and death rate of 0.5 events Myr^-1^) to be used as a fixed tree with a root age of 12.46 Myr and mean branch length of 0.89 Myr. Datasets were simulated using transition rates sampled from a truncated normal distribution (µ = 1, σ = 0.5), which were then scaled to have mean rates of 0.1, 1.0, or 10 transitions Myr^-1^ by dividing the rate matrix by the sum of the diagonal and then multiplying by the desired scalar. This resulted in a range of evolutionary rates where the expected number of transitions ranged from ∼21, 210, or 2100 transitions across the entire tree depending on the mean rate. For each transition model, 100 datasets were simulated. The transition rates of each dataset were then estimated and their maximum likelihood estimates were used to infer marginal probabilities of each character state across the tree. This procedure was repeated 10 times.

An underappreciated concern with evaluating models that differ in the number of observed states is that the probability of guessing the correct state without any additional information is simply 1/*k* states. This could, in theory, inflate the accuracy of datasets with fewer states even though the tip states themselves provide no information about the ancestral states when the rates are high (Schultz, Cocroft, & Churchill, 1996; Sober & Steel, 2011, 2014). To deal with this issue, we also calculated the mutual information, measured in bits, about ancestral states from each dataset and model (Cover & Thomas, 1991; Sober & Steel, 2011). Specifically, mutual information is a measure of how much ancestral state uncertainty is reduced by knowing the tip states (details of our derivation are given in S1). The initial uncertainty, or unconditional entropy, is set by the model – given a model of evolution and no knowledge of the extant tips, how uncertain is the best guess of the ancestral states? The remaining uncertainty after ASR, or conditional entropy, is given by the combination of the model and the tip states – given the model of evolution and knowledge of the extant tips, how uncertain is the best guess of the ancestral states? It is important to note that information, just like ancestral state reconstruction, is highly correlated with the model of evolution, and thus any results related to information will take on the assumptions of the model.

We define information as the difference between the unconditional entropy of the node states, *H*(*X*_*v*_), and the entropy of the node states conditioned on the data, *H*(*X*_*v*_ | *X*_*h*_ = *D*) (Cover & Thomas, 1991). The unconditional entropy of node *v* is defined as:

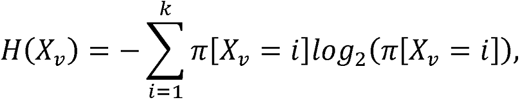

where *π* [*X*_*v*_ = *i*] is the prior probability of a node taking a particular state. For the root, the prior depends on user choice, as there are several options (Yang, Kumar, & Nei, 1995; Pagel, 1999; FitzJohn, Maddison, & Otto, 2009). Here we assume the prior probability on the root node is the expected equilibrium frequency, *π*, which is calculated directly from the transition model by solving *π**Q*** = 0. This aligns our expectation of the root node with all other internal nodes such that, in the absence of information from the tips, the probability of a particular state is assumed to be drawn from the equilibrium frequencies. In other words, the information of the tip states decreases as rates increase and, ultimately, the probability of a node state becomes completely determined by the model. We define the conditional entropy as:

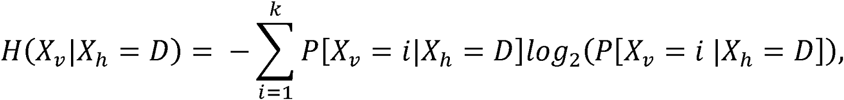

where *P*[*X*_*h*_ = *i*| *X*_*h*_ = *D*] is the conditional probability that a node is fixed as being in state *i* given the probability of observing the tip data (which is just the marginal probability of state *i*). In particular, we are interested in the average entropy of a node for all states *i* … *k*, given we observe a particular dataset, *X*_*h*_ = *D*. Thus, the conditional entropy will vary by node, but the unconditional entropy is set by the model. To produce a measure of mutual information between the observations at the tips and estimates at internal nodes, we take the difference between the conditional entropy and the unconditional entropy and average across all nodes. However, the unconditional entropies will be greater for datasets that include more states because unconditional entropy sets the upper limit of what is possible to learn. This alone could contribute to large informational differences between models with different numbers of observed states. Therefore, we also measure the proportion of maximum information gained 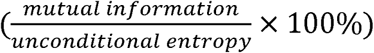.

#### 2.2.2 Evaluating hidden Markov models

We evaluated the ability to detect rate heterogeneity by simulating data under an HMM. As outlined above (see *2.1 Generalizing HMMs*), there are two major axes along which an HMM differs from standard Markov models. First, we varied the magnitude of the difference in the state-dependent process by simulating data under a model where there was: (1) no difference between the state-dependent processes (***R***_1_= ***R***_2_), (2) a 2-fold difference in rates between the state-dependent processes (e.g. if ***R***_1_’s mean rate was 1 Myr^-1^, ***R***_2_ mean rate would be 2 Myr^-1^), model in which within ***R***_1_ all transitions occur freely, but for ***R***_2_ all transition rates are zero, and (3) a 10-fold difference between the state-dependent processes, and (4) a covarion-like trait evolution is essentially “turned off” (Penny et al., 2001). For all simulation scenarios, we set the parameter-process to have equal transition rates between state-dependent processes. In addition to examining ancestral state reconstruction at nodes, we also used the new `makeSimmap` to assess how well the model captures the expected number of character changes within and among all branches in the tree. For each of the 150 datasets simulated above, we evaluated 100 simmaps per model by counting the number of transitions for a given simmap.

Next, we tested the impact of the magnitude of the asymmetry in the underlying parameter-process. We simulated data where the state-dependent process always differed by 2-fold, but for the underlying parameter-process there was: (1) no difference in transition rate (***R***_R1→R2_=***R***_R2→R1_), (2) 1.5× faster transition rate to the slower rate class (***R***_R1→R2_>*r*_R2→R1_), (3) 2× faster transition rate to the slower rate class, (4) 10× faster transition rate to the slower rate class. For each of the models described, we used simulated 500-tip phylogeny with a root age of 15.43 Myr (birth rate set to 1 event Myr^-1^, and death rate of 0.7 events Myr^-1^).

Finally, we examined how much information is available when we allow for hidden states to be observed at the tips. We used the same data generated from simulations examining state-dependent differences, but this time we did not remove the hidden state. We then fit a Markov model to this full dataset and compared it to models in which the “second character” remained unobserved.

### 2.3 Case study: reconstructing the ancestral angiosperm flower

#### 2.3.1 Background

Understanding the origin of flowering plants is widely considered to be one of the most important goals of systematic botany. In a recent study, (Sauquet et al., 2017) compiled an extensive database of floral characteristics and attempted to reconstruct the morphology of the ancestral angiosperm flower. Sauquet et al. (2017) did not present a single answer for the ancestral state because there were several possible state combinations depending on the method used and uncertainty associated with each of those estimates. Nonetheless, their hypothetical diagram of the ancestral flower as having a whorled perianth, whorled androecium, and spiral gynoecium proved controversial. For example, Sokoloff, Remizowa, Bateman, & Rudall (2018) disputed this depiction of the ancestral phyllotaxy, suggesting that some of the characters were scored incorrectly and that it seemed improbable that state combinations that are rare in the data could be the ancestral state. Sokoloff et al. (2018) instead prefer the hypothesis that the ancestral flower was either entirely whorled or entirely spiraled. In response, Sauquet et al. (2018) rescored the disputed characters and reanalyzed the dataset using the same methods as the original study. Their Bayesian analyses conformed to the predictions of Sokoloff et al. (2018), but remained highly uncertain overall. A limitation of the original study was the fact that “no comparative method exists yet to account for the potential correlation of more than two discrete characters” (Sauquet et al., 2017, but see Beaulieu & Donoghue, 2013). Given that flowers are highly integrated structures with potentially several developmental constraints the correlation between states seems a necessary prerequisite to study their evolution (Sauquet et al., 2017, 2018; Sokoloff et al., 2018). Treating the phyllotaxy of the perianth, androecium, and gynoecium as independent represents a major assumption with potentially large consequences on the ancestral state reconstruction. Indeed, Sauquet et al. (2017) conducted several pairwise correlation analyses and found that the phyllotactic patterns of these focal characters were strongly correlated.

#### 2.3.2 Worked example and methods

We limited ourselves to including only the characters related to the phyllotaxy of the perianth, androecium, and gynoecium. Although it is possible to include other characters, given the corresponding increase of parameter space, we suspect that we would not have the power to accurately infer the model and ancestral states (O’Meara et al., 2016). The dataset of Sauquet et al. (2018) has several polymorphic species as well as species for which some of the tip states are unknown. Therefore, we analyzed three separate datasets: (1) no uncertain taxa (n = 295), (2) polymorphic species included (n = 297), and (3) all taxa included (n = 780). We treated the phyllotaxy of the perianth, androecium, and gynoecium as either “whorled” or “spiral” and polymorphic species are coded to have both states. The choice dataset has major implications for model performance because corHMM will exclude state combinations that are absent from the dataset. However, the inclusion of either polymorphic or unknown states for taxa will expand state space and thus increase the number of parameters that need to be estimated. Finally, we use the C series phylogeny of Sauquet et al. (2017) in which *Amborella* is constrained as the sister to all remaining angiosperms and *Monocotyledoneae, Ceratophyllaceae*, and *Eudicotyledoneae* are constrained to form a monophyletic group.

In our case, we have three data columns each with two observed states. Because this dataset contains two or more columns of trait information, each column is automatically interpreted as an evolving character. In these cases, `corHMM` will also automatically remove dual transitions from the model since that would constitute two or more evolutionary events (Pagel, 1994; Maddison, Midford, Otto, & Oakley, 2007). However, previous work has suggested that dual transitions are possible and likely in this system (Sauquet et al. 2018; Sokoloff et al. 2018). Thus, we include both models in which dual transitions are allowed and disallowed. Our analysis without hidden states include three different model structures: `model=“ER”` (equal rates), `model=“SYM”` (symmetric rates), and `model=“ARD”` (all rates differ). The other options used (`rate.cat=1` and `nstarts=10`) specify that no hidden states are to be used and that the maximum likelihood search will be performed 10 additional times with different initial parameters.

We also include a set of analyses in which hidden states are present because it is likely that there are unobserved characters which influence the evolution of the angiosperm flower. We include four hidden state models: *ER/ER, SYM/SYM, ARD/ARD*, and *ER/ARD*. Each of these models allows for the possibility of rate heterogeneity through the inclusion of a hidden state, however the state-dependent processes differ. In the *ER/ER* model all changes between states occur at the same rate within a state-dependent process, but the magnitude of change can depend on the underlying parameter process. The *SYM/SYM* model specifies that within character changes are equally probable, but some characters may change faster than others. The *ARD/*ARD model specifies that there could be a bias towards a particular state, but this state may differ depending on whether the lineage is in ***R***_1_ or ***R***_2_. Finally, *ER/ARD* is a hybrid model which includes aspects of the equal rates model and all rates differ model.

## 3. Results

### 3.1 – Performance in Simulation

#### 3.1.1 – Increasing the number of observed characters or states

Overall, the accuracy of an ancestral state reconstruction is predominately a function of the transition rates, but there are regions of parameter space where the number of states is influential (Fig. 2a). For example, all datasets generally inferred the correct ancestral state at low rates and datasets with more states performed better at intermediate rates. However, when viewed in terms of information, datasets that contained just two states showed detectable informational loss when compared to the three- and four-state datasets. In fact, across all scenarios — low, intermediate, and especially at the highest rates — datasets with more states consistently showed more informational gain relative to the maximum information content for a given number of states (Fig. 2b). We suspect this largely reflects the impacts of homoplasy when the number of character states are restricted in the model (Sanderson & Donoghue, 1989; Steel & Penny, 2005). This is not to say that more character states are always necessary for accurate ASR. Rather, we demonstrate that there are cases when additional characters or character states enhance the accuracy of an ancestral state reconstruction and those datasets have a signal of increased information.

**Figure 2.**
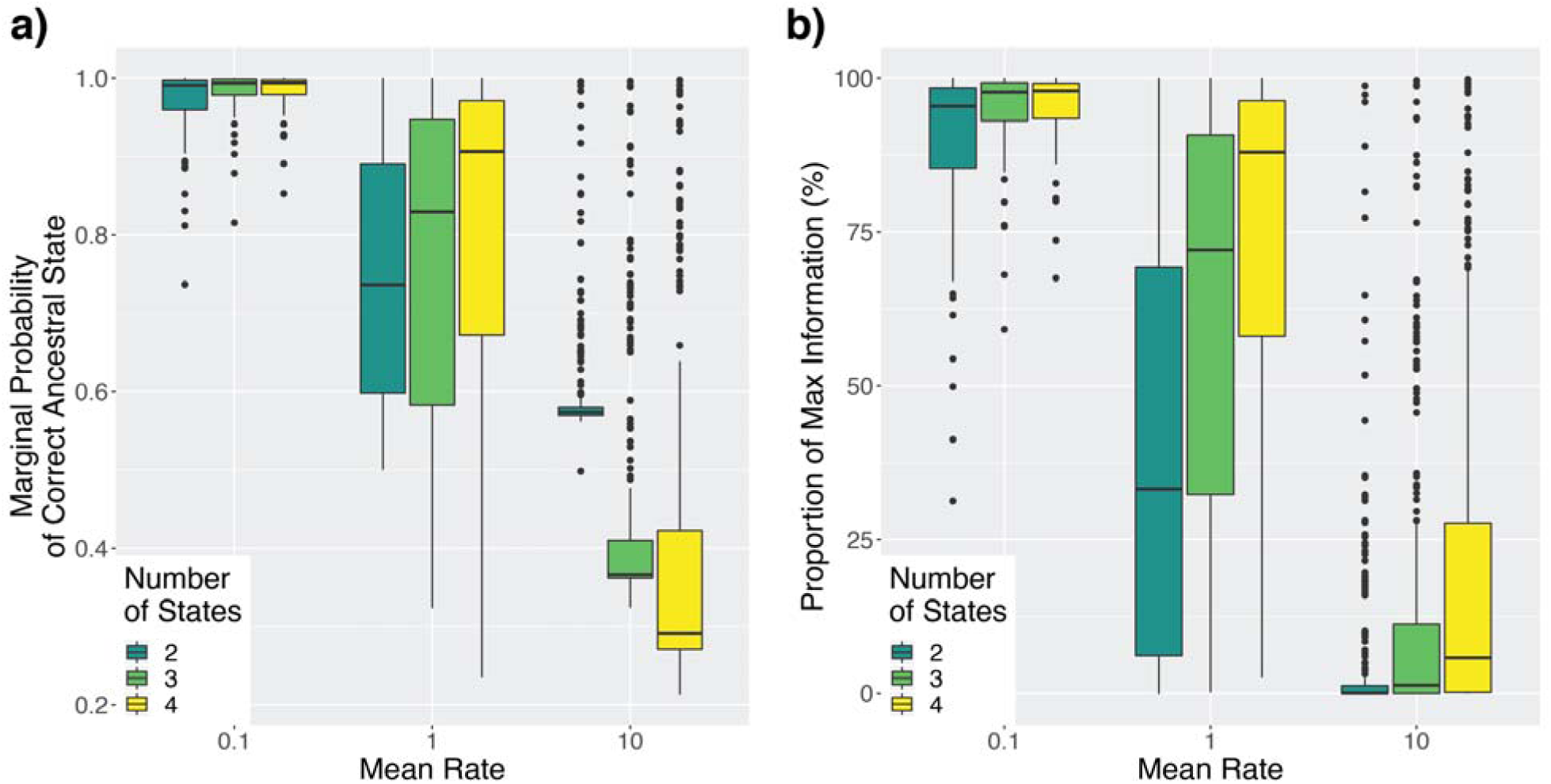
Performance of standard and hidden Markov models depending on the number of states in the dataset and mean rate. Each dataset was simulated under a mean rate of 0.1, 1, or 10 transitions Myr^-1^, with 2, 3, or 4 observed states and no hidden states. a) The marginal probability of estimating the correct ancestral state. b) The proportion of information gained about ancestral states from each dataset and model.

#### 3.1.2 – Evaluating hidden Markov models

The accuracy of ancestral state estimation, based solely on reconstructing character states at nodes, appears largely unaffected by the inclusion of hidden states regardless of differences in the state-dependent processes (Fig. 3a). However, the amount of information gained depends on both the use of an HMM and the presence of strong differences between the state-dependent processes (Fig. 3b). These seemingly contradictory results are a consequence of how we calculate information. Model uncertainty certainly comes from the increase in parameters of HMMs relative to standard Mk models and manifests in both increased model complexity and an increased number of potential ancestral states. The increase in possible ancestral states results in a higher unconditional entropy, which can actually lead to greater informational content even when the ancestral states are not known with as much certainty as a Mk model. However, as we show in section 3.1.1 increasing the number of observed states improved ancestral state reconstruction accuracy, despite a greater number of estimated parameters, and so this does not solely account for the greater ancestral state accuracy of a Mk over an HMM. We suspect it is also due to datasets fit under an HMM have added uncertainty applied to the tips because it is initially unknown which hidden state a particular taxon occupies. The greater uncertainty at the tips is likely the reason why we observe Mk models outperforming HMMs in ancestral state reconstruction, and the greater uncertainty of the model is likely why HMMs are able to extract more information from a given dataset.

**Figure 3.**
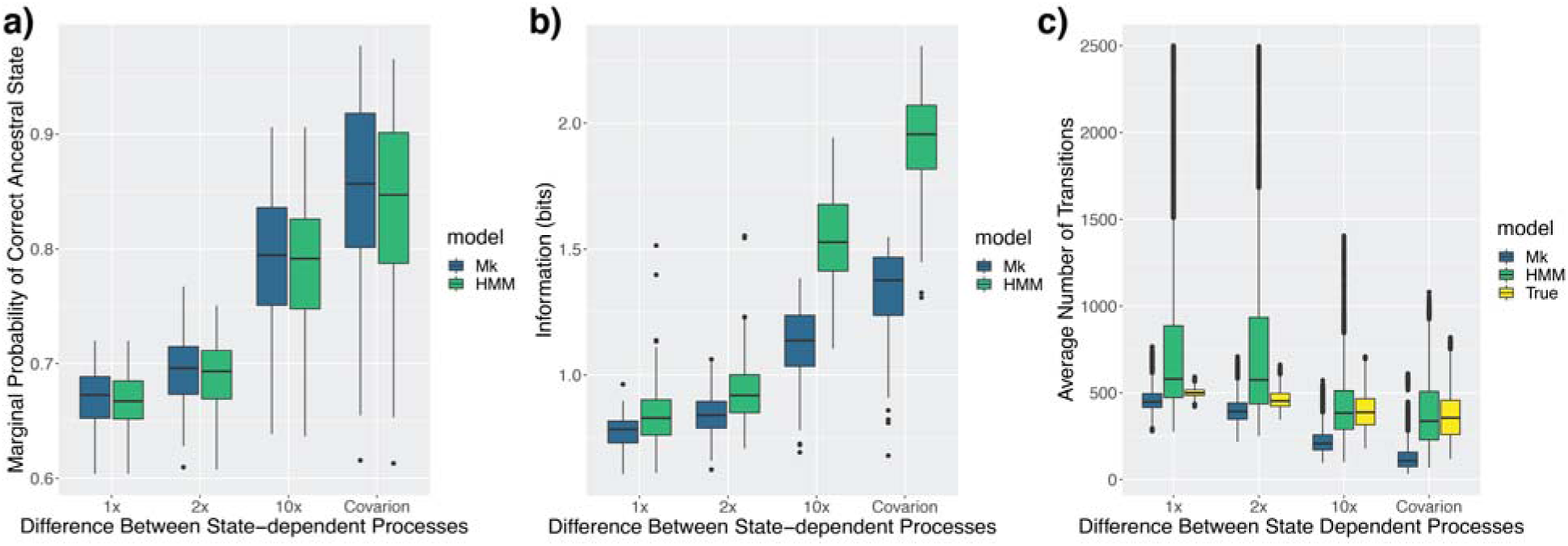
Comparison of fits from a Markov and HMM model when an HMM is the generating model. We vary the difference between state-dependent processes from no difference (1x) to complete asymmetry where state transitions occur in one state-dependent process only (i.e., “covarion” model; see *Performance in simulation*). a) The marginal probability of the correct ancestral state. b) The average amount of information (bits) for ancestral states from each dataset and model. c) The number of transitions averaged over 150 simmaps.

We found that when the generating model does not have state-dependent differences, the HMM does not pickup significant rate variation and resembles the character history implied by the standard Markov model (Fig. 4a-c). When there was no difference between the state-dependent processes, 2.6% of datasets had an AICc difference greater than 2 in support of an HMM over a Markov model. Whereas 100% of datasets supported an HMM when data was simulated under a covarion-like model. These findings suggest that HMMs are supported in datasets where rate heterogeneity is present and this can be seen qualitatively through simmap reconstructions (Fig. 4d-f). We found little effect of altering the transition rate bias of the parameter process on either ancestral state reconstruction (Fig. S1a) or information content (Fig. S1b).

**Figure 4.**
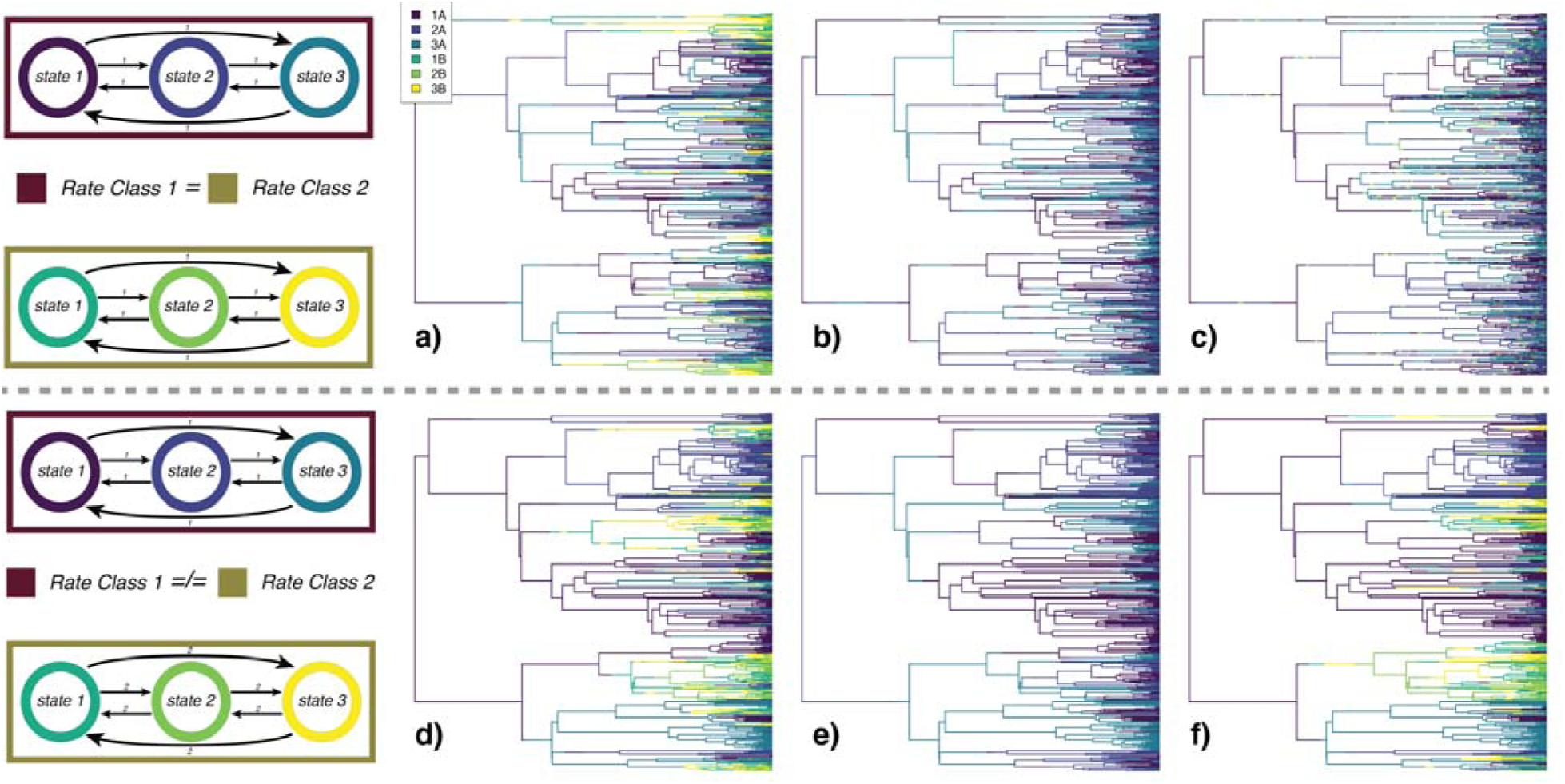
Stochastic maps demonstrating the effect of differences in the magnitude of two state-dependent processes. The first row shows data is simulated where there were no differences in the state-dependent processes, where (a) is the true generating model, (b) is one example of the character history simulated under the MLE from standard Markov model, and (c) is one example of the character history simulated under the MLE of an HMM. The second row is the same, but with data simulated with a 10-fold difference between the state-dependent processes. A Markov model does not contain a distinction between the hidden classes A and B, thus it is displayed only in terms of the states 1A, 2A, and 3A. Comparing the HMM in (c) and (f) demonstrates that an HMM will only detect a hidden state when it influences the observed, state-dependent, process.

Unsurprisingly, observing the “second character” states increased the amount of information (Fig. 5). However, as the state-dependent processes became more distinguishable, the informational gap between an HMM and including the observed second character decreased. In other words, when the evolution of an observed character changes across the phylogeny, an HMM is able to extract additional information from a dataset.

**Figure 5.**
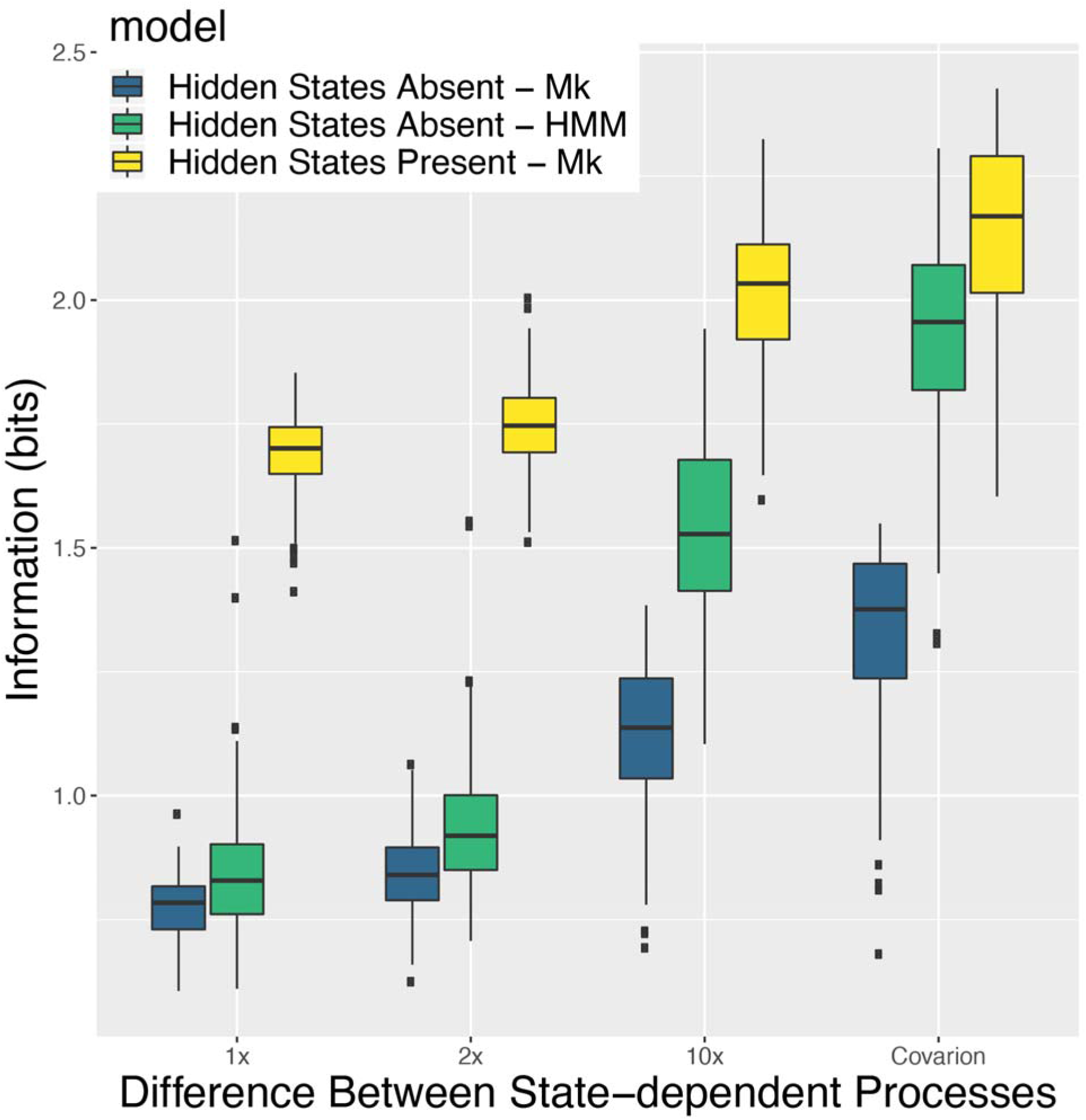
Average information when a hidden state is either directly observable or unobserved. If the hidden state is unobserved (*Hidden States Absent*), we compare the information gained when fitting a Markov model (Mk) or a hidden Markov model (HMM) to a dataset that was generated with a hidden state, but that hidden state was removed from the dataset. When the hidden state is directly observable (*Hidden States* Present) we fit a standard Mk to the full dataset that includes the potential hidden state. When the hidden state is directly observed, the datasets are comprised of 6 discrete states.

**Figure 6.**
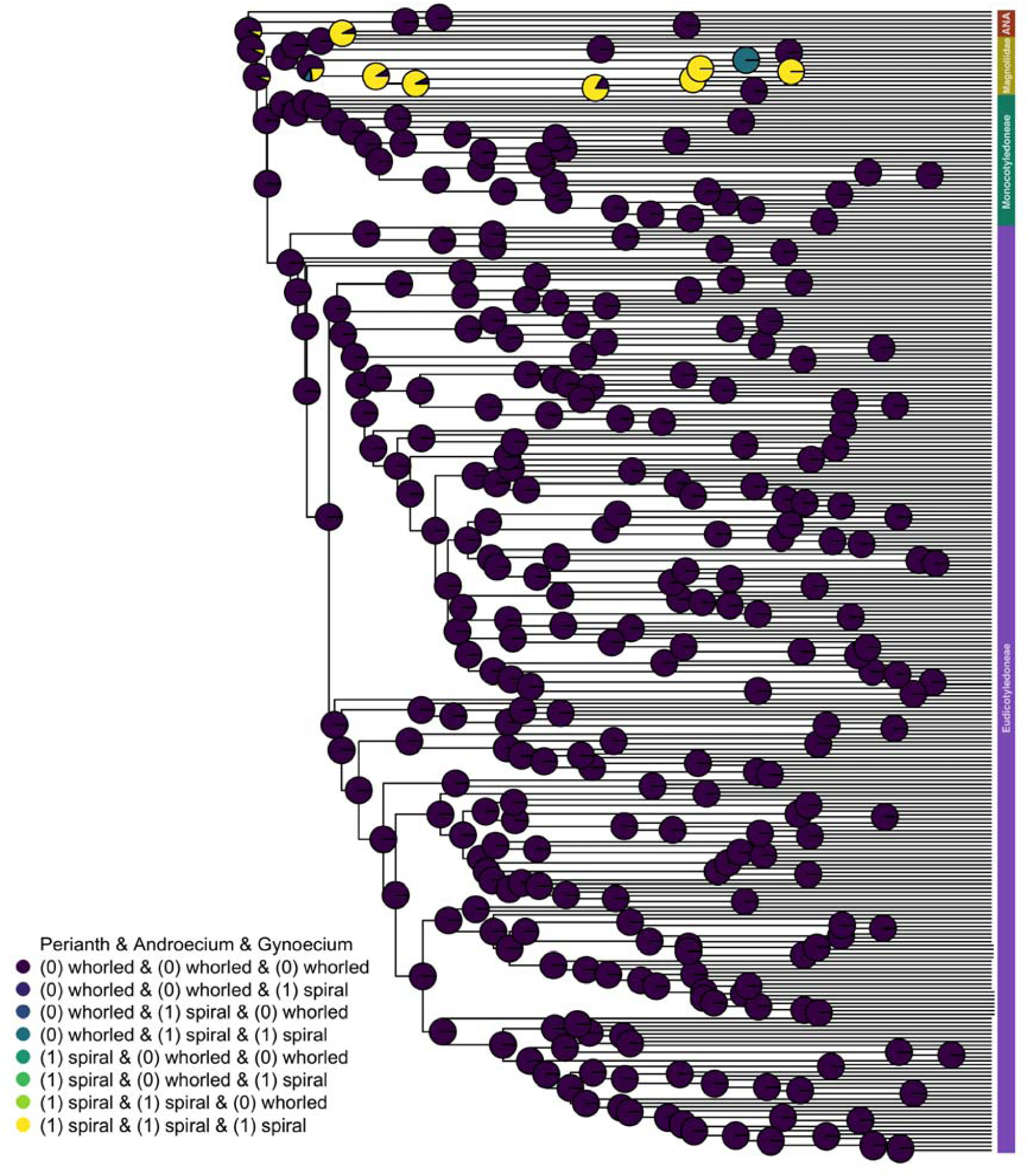
Model averaged ancestral state reconstructions of dataset two (polymorphic species are included, but species with unknown states are excluded). The marginal probability that the root state is entirely whorled is 91.7%.

### 3.2 – Case study: reconstructing the ancestral angiosperm flower

Across all three datasets our best supported model was *ER/ER*, a two rate class model with both state-dependent processes being equal rates (Table 1; Table S1). Because our modeling set included a wide range of complexity ranging from 1 estimated rate (ER) to 114 estimated rates (ARD/ARD), we used AIC weights to calculate the model averaged ancestral states for datasets individually. For all three datasets we find that an entirely whorled angiosperm flower is the most likely state (Table S2). However, we found that the preferred ancestral state is highly variable and dependent on the model and the entirely whorled angiosperm flower is likely a reflection of the *ER/ER* model’s high AIC weight within the set of tested models (Table 1; Table S1). For several of the models we found that the parameter estimates reached the upper limit of the transition rates allowed (Table S3). This could be reflection of a lack of adequate data, too many unknown and polymorphic state combinations, and/or unrealistic models included in the set. However, none of the transition rates estimated reached the upper limit for any of the best supported models.

**Table 1.** Model rankings from the maximum-likelihood analysis of the ancestral angiosperm flower for dataset two (polymorphic species included and unknown states excluded). Models separated by a “/” indicate a hidden rates model and the split distinguishes between the two state-dependent process (for example, *ER/ARD*, represents a hidden rate model where R1 is an equal rates model and R2 is an all rates differ model). Dual describe whether the model allowed for multi-state transitions (for example, if dual transitions were TRUE, then changing from entirely whorled phyllotaxy to entirely spiral phyllotaxy is allowed). AICc is the sample size corrected Aikaike Information Criterion. AICcWt is the relative likelihood of each model and is used in model averaging. Mean rate is the average transition rate for a particular model. ASR is the most likely ancestral state reconstruction for a particular model and its marginal probability. *k* rates is the number of independent rate parameters being estimated for a given model.

| Model | Dual Transitions | AICc | AICcWt | Mean Rate | ASR | *k* rates |
| --- | --- | --- | --- | --- | --- | --- |
| ER | FALSE | 208.44 | <0.01 | <0.01 | (1,R1) 93% | 1 |
| SYM | FALSE | 138.25 | <0.01 | 16.67 | (1,R1) 99% | 12 |
| ARD | FALSE | 125.53 | <0.01 | 13.56 | (5,R1) 100% | 24 |
| ER/ER | FALSE | 118.06 | 0.01 | 37.52 | (1,R2) 63% | 4 |
| SYM/SYM | FALSE | 147.3 | <0.01 | 6.25 | (1,R2) 93% | 26 |
| ER/ARD | FALSE | 132.6 | <0.01 | 4.72 | (8,R2) 87% | 27 |
| ARD/ARD | FALSE | 191.76 | <0.01 | 13.58 | (4,R1) 82% | 50 |
| ER | TRUE | 115.2 | 0.06 | <0.01 | (1,R1) 97% | 1 |
| SYM | TRUE | 193.77 | <0.01 | 5.51 | (5,R1) 100% | 50 |
| ARD | TRUE | 212.59 | <0.01 | 16.36 | (5,R1) 100% | 56 |
| **ER/ER** | **TRUE** | **109.72** | **0.93** | **<0.01** | **(1,R2) 91%** | **4** |
| SYM/SYM | TRUE | 383.59 | <0.01 | 5.47 | (8,R1) 92% | 102 |
| ER/ARD | TRUE | 220.93 | <0.01 | 3.13 | (8,R2) 93% | 59 |
| ARD/ARD | TRUE | 443.07 | <0.01 | 10.76 | (4,R1) 86% | 114 |

## 4. Discussion

Hidden Markov models are an essential tool for inferring character states across phylogenies. The new version of `corHMM`, expands the array of potential uses of HMMs by increasing the number of possible character states and allowing users to construct custom models. In addition, we demonstrated the informational advantages of using hidden Markov models versus simple Markov models. Users interested in hypothesis-driven model construction are encouraged to read through the vignette associated with the `corHMM` package. This vignette fully describes how to use the package and includes several examples of how to take a biological hypothesis and codify it into an explicit HMM.

Information theory has mainly been discussed in a theoretical context and rarely used in practice to understand empirical trait evolution (Mossel, 2003; Mossel & Peres, 2003; Townsend & Naylor, 2007; Sober & Steel, 2011, 2014; Gascuel & Steel, 2014). In this paper, we have introduced a measure for the amount of information that the tips provide the nodes during ancestral state reconstruction. Two important caveats of this measure of information. First, the data and model are taken as fixed. These are not uncommon assumptions in phylogenetic comparative methods. For example, if one is to interpret an ancestral state reconstruction it comes with the implicit assumption that the model accurately describes the evolution of the traits (Beaulieu & O’Meara, 2019). Second, mutual information, as we have defined it, only provides information relative to the specified model and specified tips. A model which is more uncertain about any ancestral state, such as an equal rates model, is likely to have a more informative ancestral state reconstruction because any deviation from an uninformed ancestral state is due to the particular values of the tip states. This does not make the equal rates model better than alternatives nor do we advocate for the use of information to assist in model selection. Instead, mutual information provides insight into the interaction between the model and tip states. Higher information of particular nodes could be indicative of an area of the phylogeny where the model’s equilibrium frequencies were different from the ancestral state reconstruction and thus the tips provided the major explanation of the ancestral state. Mutual information is also highly correlated with the rates of evolution and has the intuitive property that as rates of evolution (or time) increase the information that the tips provide to the nodes decreases (Sober & Steel, 2011).

It is important to have a biologically realistic model of trait evolution when conducting an ancestral state reconstruction. With the generalizations made to `corHMM` we have provided two distinct ways to increase the realism of phylogenetic comparative modeling. First, we have allowed for the correlated evolution of several characters and states. Whether traits are correlated because of underlying pleiotropy leading to genetic correlation (Conner et al., 2011) or selective covariance (Mahler, Revell, Glor, & Losos, 2010), at the macroevolutionary scale they are better understood in a holistic context rather than independently evolving subunits. Second, the inclusion of hidden states allows for more detailed descriptions of the evolutionary process. State-dependent processes can differ in both rate and structure and thus provide a description of heterogeneity in the tempo and mode of evolution. However, these generalizations do not exist without cost. Increased complexity of models leads to greater parameterization which can lead to poor model performance (Grundler & Rabosky, 2020). Thus, as others recommend, we suggest having multiple working hypotheses (Chamberlin, 1890; Platt, 1964; Mayr, 1997; Burnham & Anderson, 2002). The generalizations and tools available in corHMM allow for the construction of a carefully defined set of candidate models which can be compared in an information theoretic context. The Akaike Information Criterion (AIC) applies the principles of parsimony and represents a trade-off between bias and variance as a function of the dimension of the model (Forster & Sober, 1994). Combining AIC with a carefully constructed set of models leads to multi-model inference. Rather than focusing on a single best model, we can focus on the parameters from the set of our best supported models (Burnham & Anderson, 2002; Caetano, O’Meara, & Beaulieu, 2018). It is, therefore, just as important to include Mk models alongside HMMs because in cases where the increased parameterization of an HMM are unnecessary, alternative models with less parameterization are available as simpler explanations.

We have demonstrated that there is a potential informational and accuracy advantage of including additional states and characters in a simulation setting (*3.1.1 Increasing the number of observed characters or states)*. However, it remained to be seen whether modeling the correlated evolution of multiple characters would impact the ancestral state in an empirical example and whether those results match biological expectations. The controversy surrounding the phyllotaxy of the ancestral angiosperm flower is a particularly appropriate case study for the generalized version of `corHMM`, as it not only allows for the dependent evolution of several discrete characters but also includes hidden states as a fitting addition to help describe the heterogeneous evolution of angiosperms. We presented three different datasets, each allowing for a different level of polymorphism, uncertainty, and number of tip states. In our first dataset, we excluded all polymorphic species and any species with an unknown tip state. In this case, we found a two rate class *ER/ER* model was favored and the most likely ancestral state of the floral phyllotaxy was entirely whorled. This is in contrast to previous work which used a similarly constrained dataset and suggested either an ambiguous state (Maximum Parsimony result), an entirely spiral floral phyllotaxy (Maximum Likelihood result), or a spiral perianth, whorled androecium, and spiral gynoecium (reversable jump Monte Carlo Markov Chain result) (Sauquet et al. 2018 - Appendix S2, V7Csub1).

Previous work posed the question of whether we should dismiss ancestral state combinations not observed among living species (Sauquet et al. 2018). The first dataset we presented excluded any trait combinations not observed in the data. However, the other two datasets allow for an ancestral state combination that was never directly observed at the tips because these analyses include tips where the states are not completely known. In both cases, we found the hidden rates model *ER/ER* where dual transitions are allowed to be the best supported and the model averaged most likely ancestral state was an entirely whorled floral phyllotaxy. Thus, across all datasets we found that the ancestral state combination was one of the most common tip state combinations. However, this does not mean a combination of states unknown in any extant species is impossible. In fact, we find that the preferred ancestral state is highly variable and dependent on the model (Table 1; Table S1) and the entirely whorled angiosperm flower is likely a reflection of the *ER/ER* model’s high AIC weight within the set of tested models. This means that should a more realistic model be introduced in the set we could find a very different answer and highlights the importance of having a set of biologically realistic models.

## 5. Conclusion

Although there is a growing consensus that phylogenies and their associated methods are being used in ways that exceed what they can infer (Losos 2011; Maddison and FitzJohn 2015; Rabosky and Goldberg 2015; Cooper et al. 2016), we have shown that there is still under-utilized information in phylogenetic comparative datasets. First, HMMs extract signals of rate heterogeneity when it is present and, equally important, do not falsely locate signals where they are absent. Second, increased trait depth adds new information and consistently improves ancestral state reconstruction estimates. Indeed, as datasets continue to grow, so will the analytical power that biologists have for testing complex models of evolution. Finally, the inclusion of correlated trait evolution and hidden states is relevant beyond theoretical considerations and we have shown that these generalizations can change the results of an ancestral state reconstruction in empirical datasets. There is still a great deal of uncertainty in the reconstruction of the ancestral phyllotaxy of angiosperms, but by using AIC weighted marginal probabilities we have been able to take into account different biological explanations of floral evolution, eventually finding support an entirely whorled perianth, androecium, and gynoecium. Although hidden Markov models are not a perfect substitute for real observation of a hidden character, they make for a tractable and a biologically reasonable description of heterogeneity in the evolutionary process over long time scales.

## Supporting information

Modeling results

Ancestral State reconstructions

Full rate matrices

Parameter process figure

## Availability

This open source software is written entirely in the R language and is freely available through the Comprehensive R Archive Network (CRAN) at: https://cran.r-project.org/web/packages/corHMM/index.html.

The scripts used, simulation results, model fits, and case study data are available at: https://datadryad.org/stash/dataset/doi:10.5061/dryad.vx0k6djpg.

## Acknowledgements

We thank Hervé Sauquet and an anonymous reviewer for their insightful comments on an earlier version of this paper. We also thank members of the Beaulieu lab and colleagues at the University of Arkansas for their comments and for general discussions of the ideas presented here. Finally, we would like to specifically thank Andrew Alverson, Brian O’Meara, and Adam Siepielski for their insightful critiques and helpful edits on an earlier version of this manuscript.

## Author Contributions

J.D.B. and J.M.B. designed research; J.D.B performed research and analyzed data; and J.D.B. and J.M.B. wrote the paper.

