## Supplementary material for "Generalized Hidden Markov Models for Phylogenetic Comparative Datasets": Parameter process figure

Supplemental Material 1

***Mutual Information***

We define information as the difference between the unconditional entropy of the node states, $H\left( X_{v} \right)$, and the entropy of the node states conditioned on the data, $H\left( X_{v}|X_{h}=D \right)$ (Cover & Thomas, 1991). The unconditional entropy of node $v$ is defined as:

$$H\left( X_{v} \right)=-\sum_{i=1}^{k} \pi\left[ X_{v}=i \right]{log}_{2}\left( \pi\left[ X_{v}=i \right] \right),$$

where $\pi\left[ X_{v}=i \right]$ is the prior probability of a node taking a particular state. For the root, the prior depends on user choice, as there are several options (Yang, Kumar, & Nei, 1995; Pagel, 1999; FitzJohn, Maddison, & Otto, 2009). Here we assume the prior probability on the root node is the expected equilibrium frequency, $\pi$, which is calculated directly from the transition model by solving $\pi\boldsymbol{Q}=0$. This aligns our expectation of the root node with all other internal nodes such that, in the absence of information from the tips, the probability of a particular state is assumed to be drawn from the equilibrium frequencies. In other words, the information of the tip states decreases as rates increase and, ultimately, the probability of a node state becomes completely determined by the model. We define the conditional entropy as:

$$H\left( X_{v}|X_{h}=D \right)= -\sum_{i=1}^{k} P\left[ X_{v}=i|X_{h}=D \right]{log}_{2}\left( P\left[ X_{v}=i |X_{h}=D \right] \right),$$

where $P\left[ X_{v}=i|X_{h}=D \right]$ is the conditional probability that a node is fixed as being in state $i$ given the probability of observing the tip data (which is just the marginal probability of state *i*). In particular, we are interested in the average entropy of a node for all states$i\ldots k$, given we observe a particular dataset, $X_{h}=D.$ Thus, the conditional entropy will vary by node, but the unconditional entropy is set by the model. To produce a measure of mutual information between the observations at the tips and estimates at internal nodes, we take the difference between the conditional entropy and the unconditional entropy and average across all nodes. However, the unconditional entropies will be greater for datasets that include more states because unconditional entropy sets the upper limit of what is possible to learn. This alone could contribute to large informational differences between models with different numbers of observed states. Therefore, we also measure the proportion of maximum information gained $(\frac{mutual information}{unconditional entropy}\times100\%)$.

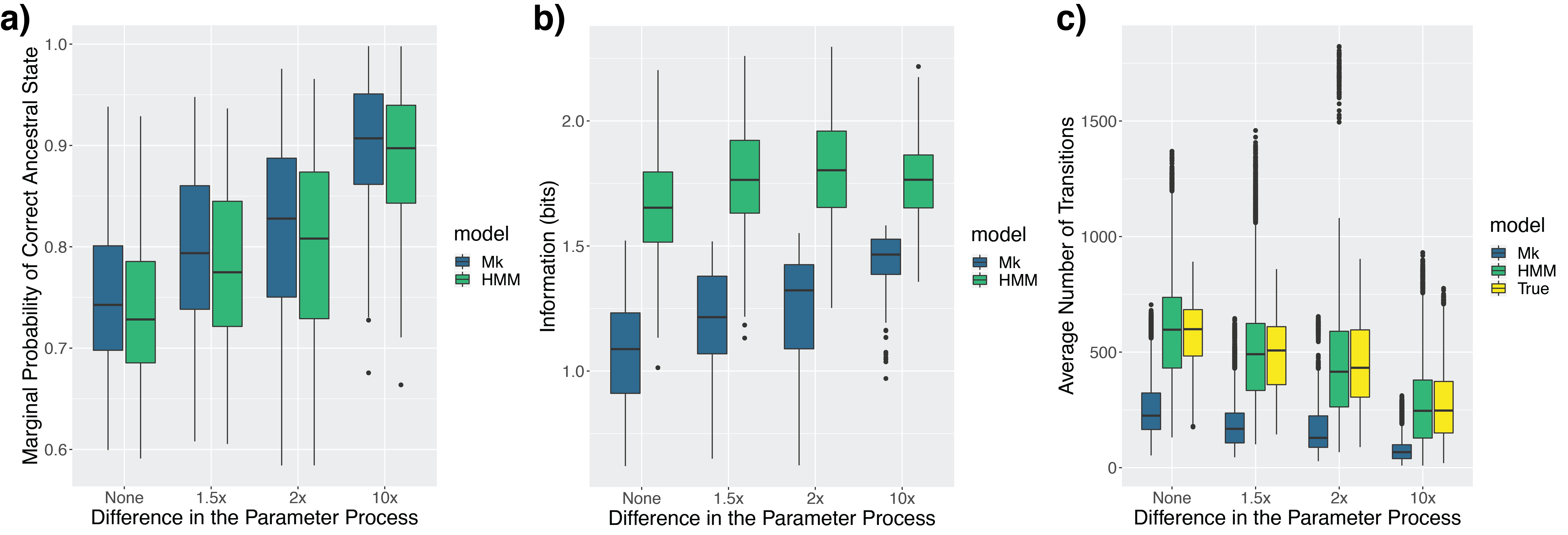

**Figure S1**. Comparison of a standard Markov and HMM model when an HMM is the generating model. We vary the bias from rate class 1 to rate class 2 from no difference (none) to a 10-fold difference (10x). a) The marginal probability of the correct ancestral state. b) The average amount of information the tips provide the nodes. c) The number of transitions averaged over 150 simmaps.
